# Introducing *π*-HelixNovo for practical large-scale de novo peptide sequencing

**DOI:** 10.1101/2023.07.15.549133

**Authors:** Tingpeng Yang, Tianze Ling, Boyan Sun, Zhendong Liang, Fan Xu, Xiansong Huang, Linhai Xie, Yonghong He, Leyuan Li, Fuchu He, Yu Wang, Cheng Chang

**Affiliations:** Peng Cheng Laboratory, Shenzhen, 518055, China; State Key Laboratory of Proteomics, Beijing Proteome Research Center, National Center for Protein Sciences (Beijing), Beijing Institute of Lifeomics, Beijing, 102206, China; Tsinghua Shenzhen International Graduate School, Shenzhen, 518055, China; School of Life Sciences, Tsinghua University, Beijing, 100084, China; Research Unit of Proteomics Driven Cancer Precision Medicine, Chinese Academy of Medical Sciences, Beijing, 102206, China

## Abstract

De novo peptide sequencing is a promising approach for novel peptide discovery. We use a novel concept of complementary spectra to enhance ion information and propose a de novo sequencing model *π*-HelixNovo based on Transformer architecture. *π*-HelixNovo outperforms other state-of-the-art models and enhances the taxonomic resolution of gut metaproteome, taking a significant step forward in de novo sequencing.

## Main

Tandem mass spectrometry (MS), as the mainstream high-throughput technique to identify protein sequences, plays an essential role in proteomics research by generating mass spectra (MS1, MS2) and then analysing the corresponding peptide sequences^[1]^. In a routine proteomics experiment process (**Supplementary Fig. 1a**), proteins are first digested by protease, producing a mixture of various peptides. The peptides are then separated using liquid chromatography and analysed by MS, which produces MS1 spectra, displaying the mass and charge information of the peptides. In the data-dependent acquisition (DDA) mode, each peptide is then subjected to a fragmentation operation in the mass spectrometer, and its MS2 spectrum is generated. For peptide sequence identification, an MS2 spectrum and its precursor information (i.e., the precursor mass and charge) are used to reconstruct the corresponding peptide sequence. By identifying all the possible peptides produced from the digestion of a specific protein, the entire protein sequence can be reconstructed (**Supplementary Fig. 1b**). In the process of fragmenting peptides, collision-induced dissociation (CID)^[2]^ and higher-energy C-trap dissociation (HCD)^[3]^ are often used to cleave the peptide bond between amino acids. In theory, if the collision energy is controlled in such a way that only the peptide bonds are broken and no other chemical bonds are broken during the collision, two residues known as b and y ions^[4]^ will be produced by each peptide segment during the collision. After multiple collisions, all the residues, at least in theory, will be displayed on the MS2 spectrum (the ideal spectrum, **Fig. 1a**). Using an ideal MS2 spectrum, peptide sequence identification is straightforward. However, in a typical experimental MS2 spectrum, a series of additional signals may be generated because of the unexpected breaking of other chemical bonds and the combination of ions during the collision process^[6]^. It is crucial to note that the additional signals can be classified into two main types (**Fig. 1a**). The first type consists of low-intensity peaks that can be considered Gaussian noise. The second type composes higher-intensity peaks that are usually formed by the uncommon a-ions^[4]^ resulting from peptide fragmentation and the combination of various ions. These peaks strongly correlate with specific peptide sequences, making them unique and useful signals to identify corresponding peptides. Furthermore, certain b and y ion signals may be missing^[7]^ in the MS2 spectrum, and the intensity equality of a pair of b and y ions may also be disrupted.

**Fig. 1.**
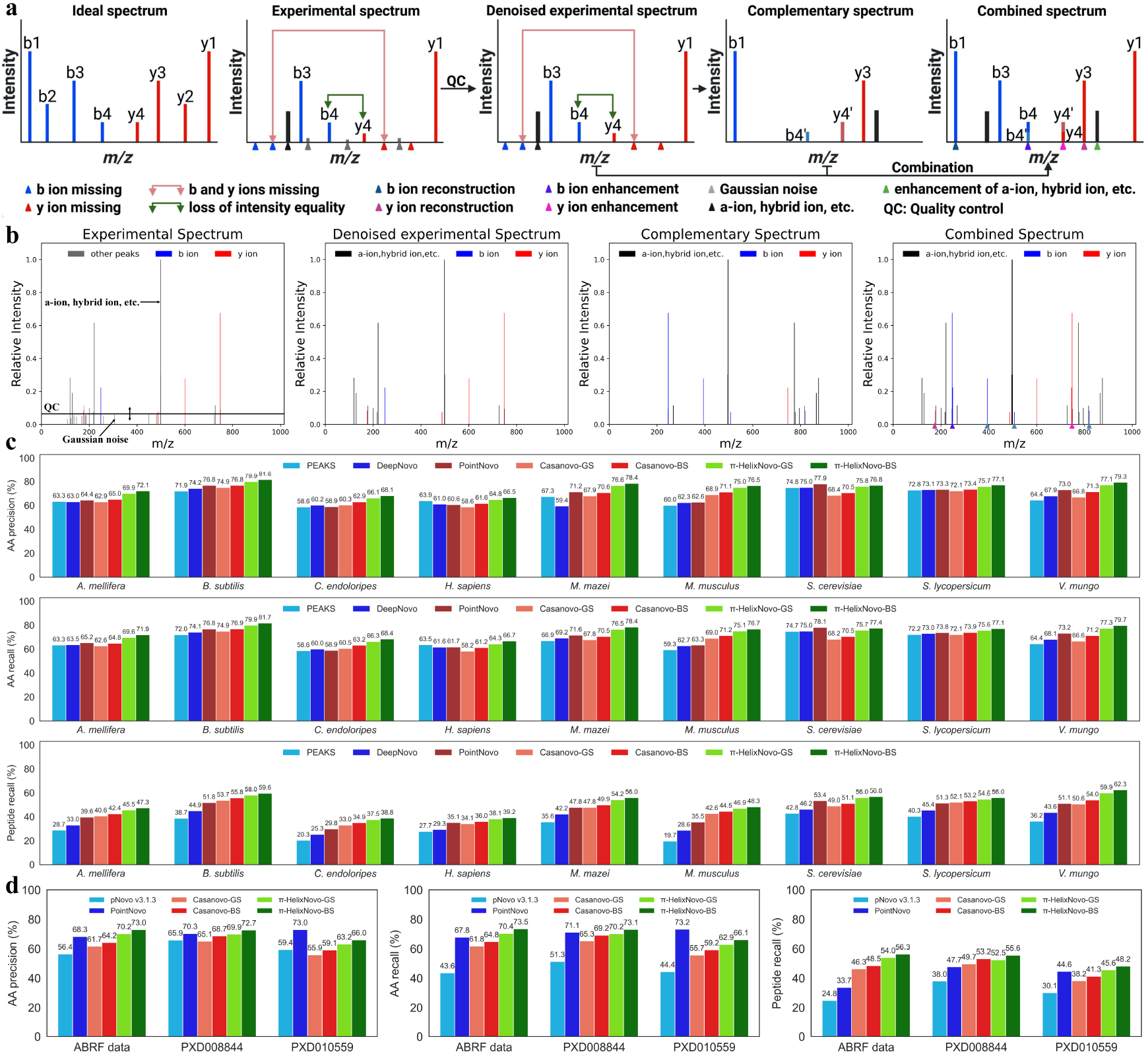
The overview of the complementary spectrum and the comparative experiments between *π*-HelixNovo and other models such as PEAKS, DeepNovo, pNovo, PointNovo, and Casanovo. (a) The difference between the ideal and experimental spectrum, the denoising of the experimental spectrum, and the complementary and combined spectrum definitions. Hybrid ions represent the combination of various ions. (b) A real example to verify the complementary spectrum’s effectiveness at enhancing the experimental spectrum. Note that the b and y ions are labelled using the PROSPECT^[5]^ dataset, and generally, the labels are not available in practice. (c) The AA precision, AA recall, and peptide recall for *π*-HelixNovo trained on the nine-species benchmark dataset using the leave-one-out cross-validation framework, compared to PEAKS, DeepNovo, PointNovo, and Casanovo. (d) The AA precision, AA recall, and peptide recall for *π*-HelixNovo trained on the merged dataset, compared to pNovo, PointNovo, and Casanovo. Note that GS and BS represent two kinds of search strategies: greedy search and beam search (*n*_*beams*_ *=* 2), respectively.

To address the above issues, we introduce a complementary spectrum as a supplement to the experimental spectrum, which has been proven useful in database search methods ^[8]^. MS2 data can be represented as a set of peaks {(*m*_1_/*z*_1_, *I*_1_*m*), ⋯ (*m*_*N*_/*z*_*N*_, *I*_*N*_)}, where *N* is the number of peaks, and *m*_*i*_, *z*_*i*_ are the mass and charge of the i-th peak. Inspired by the fact that the sum of the masses of a pair of b and y ions equals the mass of the peptide, and the intensity of a pair of b and y ions is theoretically equal, the complementary spectrum is defined as: {((*m*_*pre*_ – *m*_1_)/*z*_1_, *I*_1_), ⋯ ((*m*_*pre*_ – *m*_*N*_)/*z*_*N*_, *I*_*N*_)}, where *m*_*pre*_ is the mass of the peptide precursor. As shown in **Fig. 1a**, the complementary spectrum can reconstruct some b and y ions when some pairs of b and y ions are incomplete, enhancing the existing complete pairs of b and y ions. It is important to note that if a pair of b and y ions is enhanced, there may be a noticeable intensity inequality compared to the original pair (the intensity equality loss between b ion and y ion in the same pair). Additionally, it is worth noting that the complementary spectrum is generated after an experimental spectrum has undergone a quality control process that removes most of the Gaussian noises. This approach has the potential to increase the signal-to-noise ratio of any experimental spectrum in which some of the b and y ions are missing. **Fig. 1d** illustrates how adding the corresponding complementary spectrum can enhance an experimental spectrum.

*π*-HelixNovo is a de novo sequencing model that utilises Transformer architecture. The model comprises two main components: the Encoder and the Decoder. The Encoder component encodes the experimental spectrum using Peak Encoder E and the complementary spectrum using Peak Encoder C. The encoding results of Peak Encoder E and Peak Encoder C are concatenated and passed on to the Transformer Encoder. In the Decoder component, the Precursor Encoder processes precursor information and transmits it to the Transformer Decoder at the beginning of decoding. The first amino acid of the peptide is generated by searching the outputs of the Linear and Softmax modules. Subsequently, the Amino acid encoder (AA Encoder) and position encoder processes the previously generated amino acid, and the Transformer Decoder generates the following amino acid. This process continues until the end symbol is output or the maximum inference length is reached, thereby completing the de novo sequencing. Refer to **Supplementary Fig. 2** for a graphical representation of the model.

A series of comparative experiments have been conducted to compare *π* -HelixNovo’s performance with existing state-of-the-art de novo sequencing models (i.e., pNovo^[9]^, PEAKS^[10]^, DeepNovo^[11]^, PointNovo^[12]^, and Casanovo^[13]^). We first evaluated the complementary spectrum’s ability to enhance model performance in the same way as other de novo sequencing models by using three evaluation metrics: amino acid precision (AA precision), amino acid recall (AA recall), and peptide recall. We developed three kinds of Peak Encoder C, namely Encoder-1, Encoder-2, and Encoder-3, to encode the complementary spectrum. Models using these different encoders for the complementary spectrum were trained on the nine-species benchmark dataset (*A. mellifera, B. subtilis, C. endoloripes, H. sapiens, M. mazei, M. musculus, S. cerevisiae, S. lycopersicum*, and *V. mungo*) with the *V. mungo* dataset as the test dataset and the remaining datasets as the training dataset and compared with Casanovo^[13]^ using two different search strategies. As shown in **Supplementary Fig. 3**, utilising Encoder-1 (same as Peak Encoder E), Encoder-2, or Encoder-3 (*π*-HelixNovo) to encode the complementary spectrum led to improvements in de novo sequencing model performance as compared to Casanovo. This finding further demonstrates that the complementary spectrum can effectively enhance the performance of de novo sequencing deep learning models. It is worth noting that Encoder-3 (*π*-HelixNovo) performed best, confirming its selection as the peak encoder for the complementary spectrum in *π*-HelixNovo.

Due to the high computational cost of beam search, *π*-HelixNovo utilises greedy search as its initial search strategy. **Fig. 1c** illustrates the results of experiments on the nine-species benchmark dataset using the leave-one-out cross-validation framework and the greedy search strategy. In terms of peptide recall, *π*-HelixNovo-GS (greedy search) consistently outperformed PEAKS, DeepNovo, PointNovo, and Casanovo-GS. In terms of peptide recall, *π*-HelixNovo-GS outperformed the other four models by the following percentages (compared with the best performance in the other four models) in each cross-validation experiment: *A. mellifera* (+4.9%), *B. subtilis* (+4.3%), *C. endoloripes* (+4.5%), *H. sapiens* (+3.0%), *M. mazei* (+6.4%), *M. musculus* (+4.3%), *S. cerevisiae* (+2.6%), *S. lycopersicum* (+2.5%), and *V. mungo* (+8.8%). In terms of AA precision and AA recall, *π*-HelixNovo-GS also outperforms all other models. Although PointNovo achieved better results than *π*-HelixNovo-GS in AA precision and AA recall for the *S. cerevisiae* dataset, it still did not surpass *π* -HelixNovo-GS in peptide recall, the most crucial evaluation metric for practical applications. This is mainly due to PointNovo’s high (but not equal to 100%) amino acid prediction accuracy in some peptide sequences, resulting in a phenomenon of high overall amino acid accuracy but low peptide recall. As a result, *π* -HelixNovo-GS has consistently demonstrated improved performance in the experiments described above. Upon evaluating the models trained on the merged dataset, similar results were observed. **Fig. 1d** illustrates that *π*-HelixNovo-GS’s peptide recall surpasses the maximum values of the other three models’ peptide recall (pNovo v3.1.3, PointNovo, and Casanovo-GS), with an increase in percentage for each test dataset: ABRF data (+7.7%), PXD008844 (+2.8%), and PXD010559 (+1.0%).

To further enhance the performance of *π*-HelixNovo, beam search was employed as the search strategy. As shown in **Supplementary Fig. 4** and **Supplementary Fig. 5**, evaluation metrics were tested under different numbers of beams (*n*_*beams*_), and the results indicated that the main improvement occurs when *n*_*beams*_ transitions from 1 to 2. We also evaluated the time cost of *π*-HelixNovo under different *n*_*beams*_ on an NVIDIA Tesla V100 with 32GB of memory, and it showed that as *n*_*beams*_ increases, the time cost also increases rapidly (**Supplementary Table 1**). Therefore, the value of *n*_*beams*_ was ultimately determined to be 2, balancing performance and time cost. **Fig. 1** also displays the performance of *π*-HelixNovo-BS (beam search) and Casanovo-BS with *n*_*beams*_ set to 2, and extra improvements were gained as expected. Specifically, *π* - HelixNovo-BS exceeds the maximum values of the other four models (PEAKS, DeepNovo, PointNovo, and Casanovo-BS) in terms of peptide recall by the following percentages for each cross-validation experiment: *A. mellifera* (+4.9%), *B. subtilis* (+3.8%), *C. endoloripes* (+3.9%), *H. sapiens* (+3.2%), *M. mazei* (+6.1%), *M. musculus* (+3.8%), *S. cerevisiae* (+3.4%), *S. lycopersicum* (+2.8%), and *V. mungo* (+8.3%). Furthermore, when evaluating the models trained on the merged dataset, *π*-HelixNovo-BS’s peptide recall surpassed the maximum values of the other three models (pNovo v3.1.3, PointNovo, and Casanovo-BS), with an increase in percentage for each test dataset: ABRF data (+7.8%), PXD008844 (+2.4%), and PXD010559 (+3.6%). It’s worth noting that, to ensure fairness in the experiments above, we also used beam search as the search strategy when evaluating Casanovo (Casanovo-BS). The results provide additional evidence of *π*-HelixNovo’s exceptional performance and the complementary spectrum’s effectiveness in enhancing de novo sequencing accuracy.

We also assessed the performance of *π* -HelixNovo on peptides with varying lengths, as illustrated in **Supplementary Fig. 6** and **Supplementary Fig. 7**. In all instances, *π*-HelixNovo outperformed Casanovo (*π*-HelixNovo-BS > *π*-HelixNovo-GS > Casanovo-BS > Casanovo-GS). Notably, certain results contain singular values, which are mainly due to the small number of test samples resulting in statistically insignificant outcomes. For example, in the *H. sapiens* dataset test, the model’s performance fluctuates significantly when the peptide length is 5 or 6. Additionally, when the peptide segment length exceeds 30, the statistical results fluctuate due to the limited sample size. Overall, the results demonstrate the stable performance improvement of *π*-HelixNovo compared to other de novo sequencing models.

Next, we trained *π*-HelixNovo on the MSV000081142 dataset to achieve better performance, which contains about 4.3 million PSMs. We thought that training PandoNovo with more PSMs would achieve better performance. The model was evaluated on three test datasets, as described in **Fig. 1d**. To make the comparison fair, we ensured that the MSV000081142 dataset and the three test datasets did not share any PSMs. As shown in **Supplementary Table 2**, *π*-HelixNovo’s evaluation metrics improved greatly for each test dataset.

Furthermore, we evaluated *π*-HelixNovo trained on the MSV000081142 dataset’s ability to analyse PSMs with never-before-seen peptide sequences and made a comparison with *π* - HelixNovo trained on the merged dataset. Specifically, for each test set, we screened out PSMs with a peptide sequence that occurs in the training datasets of MSV000081142 or merged datasets. **Supplementary Fig. 8** shows the performance of two *π*-HelixNovo models under different search strategies, and it’s evident that *π*-HelixNovo performs better when trained on larger datasets. Furthermore, it should be emphasised that *π*-HelixNovo trained on the MSV000081142 dataset achieves an average performance of 53.5% in terms of peptide recall on three never-before-seen test datasets, which substantially advances the field of de novo sequencing.

Finally, we evaluated the performance of *π*-HelixNovo in metaproteomics applications where peptide precision and taxonomic resolution are of crucial importance. In metaproteomics, due to high sample complexity, taxonomic annotation of peptides relies on their uniqueness across a large database such as all UniProt entries^[14]^. An improved ability to increase the number of taxon-unique peptides would imply an advancement of the approach. Using a gut metaproteomic dataset from germ-free mice gavaged with a microbiome consortium comprising 17 defined human gut commensal species^[15]^, our results showed that *π* -HelixNovo outperformed Casanovo by identifying a greater number of novel peptides with improved taxonomic resolutions. As shown in **Fig. 2a**, in a 5-fold cross-validation experiment, *π*-HelixNovo outperformed Casanovo with the following percentage improvements in AA precision, AA recall, and peptide precision using greedy search strategy: 6.9%, 6.9%, and 7.0%, respectively. With the beam search strategy, *π*-HelixNovo achieves improvements of 5.8% in AA precision, 5.8% in AA recall, and 5.7% in peptide precision. Next, we performed de novo sequencing of those MS2 spectra that remained unidentified by database search. *π*-HelixNovo generated a greater number of peptide species (2588 vs 2224) with high confidence using the quality control T\D\DS. Notably, the overall peptide counts for both models were similar (**Fig. 2b**). Moreover, *π*-HelixNovo uncovered a larger number of bacterial-specific peptides (586 vs 350) and species-specific peptides (63 vs 15), highlighting the higher taxonomic resolution achieved by *π*-HelixNovo (**Fig. 2c-d, Supplementary Table 3**). Specifically, *π*-HelixNovo demonstrated a higher peptide precision reflected by the discovery of correct bacterial species matches (11 vs 7) and a lower occurrence of incorrect bacterial lineages (1 on the species level vs 2 on genus and phylum level). Overall, *π*-HelixNovo generated a greater number of peptide species than Casanovo (total: 33,321 vs 28,694, specific: 11,169 vs 6,542) with high confidence using the quality control T\D (**Fig. 2e**). These correspond to higher taxon-unique peptides identified by *π*-HelixNovo than by Casanovo: phylum (16,555 vs 13,928, +18.9%), genus (8,794 vs 7,335, +19.9%), and species (3,516 vs 2,845, +23.6%), respectively (**Fig. 2f**), evidencing improved taxonomic resolution in *π*-HelixNovo-based analysis.

**Fig. 2.**
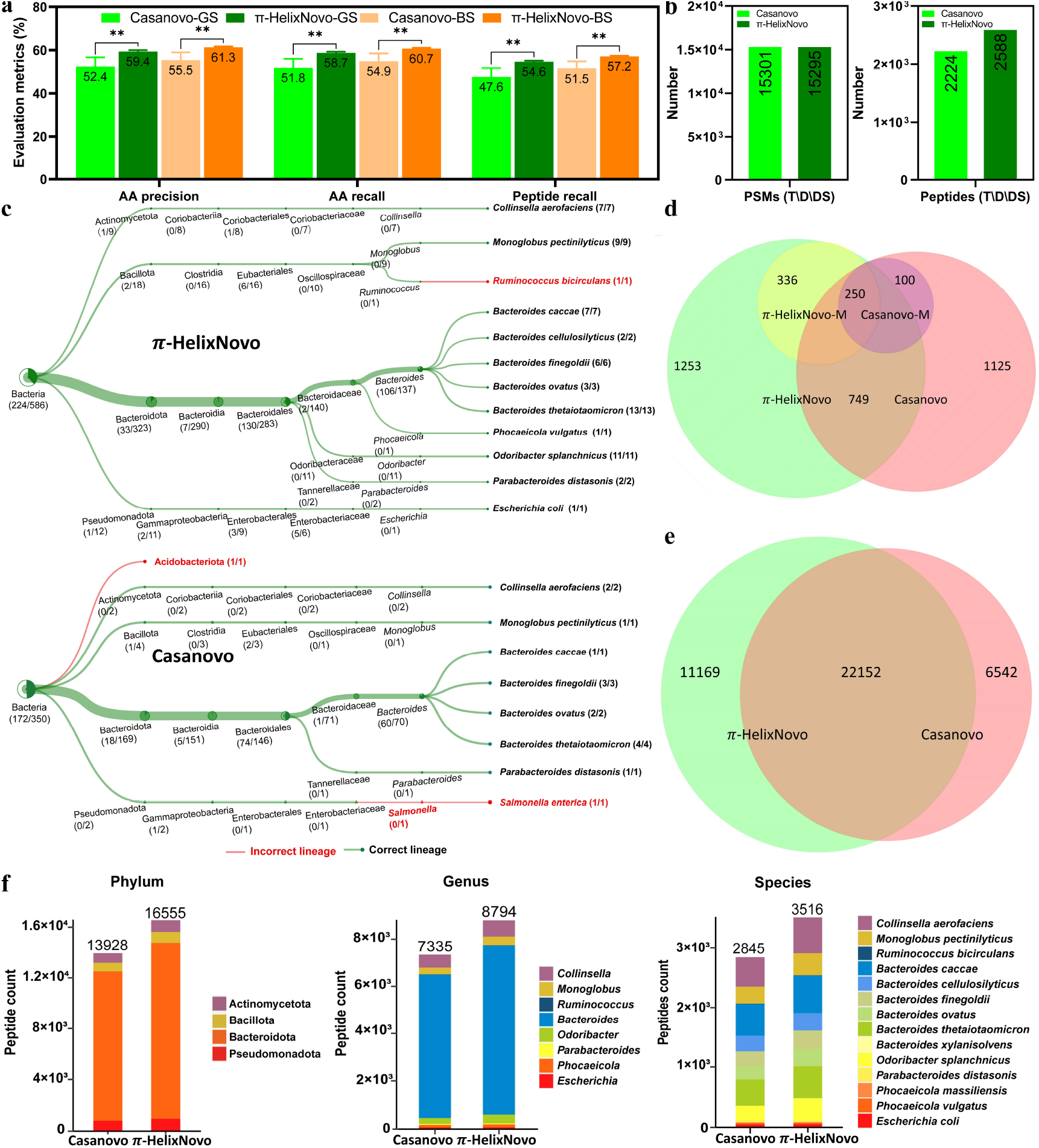
The advantages of *π*-HelixNovo in metaproteomics application. (a) The 5-fold cross-validation on PSMs identified by database search for *π*-HelixNovo, compared with Casanovo. Data are shown as mean ± SD. *p < 0.05, **p < 0.01, ***p < 0.001, ****p < 0.0001, student’s t-test. (b) The analyses of PSMs and peptides through the quality control T\D\DS: in target database but absent in decoy database and database search results. Note that these results are obtained from de novo sequencing on MS2 that cannot be identified by database search. (c) The species-level identification analysis of peptides obtained from (b). (d) The Venn diagram of peptides after quality control T\D\DS (*π* -HelixNovo, Casanovo) and the microbial-specific peptides (*π* -HelixNovo-M, Casanovo-M). (e) The Venn diagram of peptides after quality control T\D: in target database but absent in decoy database. (f) The analyses of peptides (T\D) at phylum, genus, and species levels. Note that taxa identified by less than 3 unique peptides are not included in the stacked column bar plots.

In summary, we propose the use of *π* -HelixNovo with the support of the complementary spectrum and have demonstrated that *π*-HelixNovo achieves significantly more accurate results than other state-of-the-art methods in a series of comparative experiments. We also trained *π*-HelixNovo on a larger dataset to further improve *π*-HelixNovo’s performance, and *π*-HelixNovo showed good analytical ability for MS2 of never-before-seen peptide sequences. Finally, we have utilised *π*-HelixNovo to identify previously unseen peptides in a consortium of gut microbiome species where *π*-HelixNovo also demonstrated superior performance. *π*-HelixNovo offers fresh insights into de novo peptide sequencing and advances it significantly towards practical applications.

## Methods

### Data sets

A nine-species benchmark dataset was utilised to assess the effectiveness of *π*-HelixNovo and compare it to other de novo sequencing models (PEAKS, DeepNovo, PointNovo, and Casanovo). The dataset comprised approximately 1.5 million peptide-spectrum matches (PSMs) from nine distinct species, and a leave-one-out cross-validation framework was employed in the same way as previous work. The model was trained on eight species and then tested on the remaining one. Consequently, nine experiments were carried out to evaluate the performance of *π*-HelixNovo.

Additionally, a merged dataset (approximately 0.6 million PSMs) of four high-resolution MS/MS datasets: PXD008808, PXD011246, PXD012645, and PXD012979, three other high-resolution MS/MS datasets: ABRF, PXD008844, PXD010559 were used to compare the performance of *π*-HelixNovo with pNovo, PointNovo, and Casanovo. Models were trained on the training set of the merged dataset, verified on the validation dataset of the merged dataset, and tested on the test datasets (ABRF, PXD008844, and PXD010559), which is in agreement with the testing procedure of pNovo and PointNovo.

The MSV000081142 dataset comprised around 4.3 million PSMs, and it was partitioned into training and validation sets at a ratio of 99.9:0.1 for training a more powerful *π*-HelixNovo model. To facilitate the comparison, ABRF, PXD008844, and PXD010559 were used as the test sets.

The MSV000082287 dataset contained MS2 data of human gut bacteria proteins in which approximately 26.5 million PSMs were identified through database search and 9.5 million MS2 remained unidentified^[15]^. The database was used to evaluate the performance of *π*-HelixNovo in identifying peptides of gut microbes, compared to Casanovo. Specifically, the PSMs were partitioned into training and validation sets, *π*-HelixNovo and Casanovo were then trained on the training set and evaluated on the validation set using a 5-fold cross-validation method. Finally, de novo sequencing was performed on the unidentified MS2 using *π*-HelixNovo and Casanovo.

### Quality control for the experimental spectrum

The quality control process involved the following steps:

1. Removing peaks with an *m*/*z* value outside of the [(*m*/*z*)_*min*_, (*m*/*z*)_*max*_] range.
2. Removing peaks within the given mass tolerance in Dalton around the precursor mass.
3. Removing peaks with an intensity value lower than a threshold value *I*_*c*_.
4. Selecting the top *n*_*p*_ peaks by arranging them in descending order of intensity after the above three steps have been performed.

where (*m*/*z*)_*min*_, (*m*/*z*)_*max*_, *I*_*c*_ are specified as 50.52564895, 2500.0, and 0.01 respectively.

### Details of the *π*-HelixNovo model

#### Peak Encoder E and C

Each *m*/*z* value in the experimental spectrum is encoded as a d-dimensional vector using a fixed sinusoidal embedding. The j-th dimension of the i-th *m*/*z* value (*f*_*i,j*_) is calculated as follows:

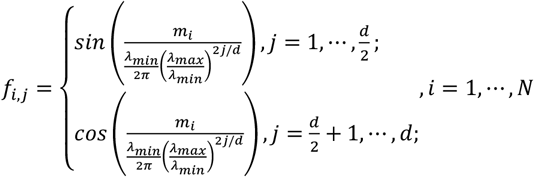

where *λ*_*min*_ *=* 0.001, *λ*_*max*_ *=* 10,000, while *d* represents the dimension of the Transformer layer for *π*-HelixNovo (*d*_*m*_). For the complementary spectrum, a similar calculation is used to obtain the j-th dimension of the i-th *m*/*z* value 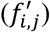:

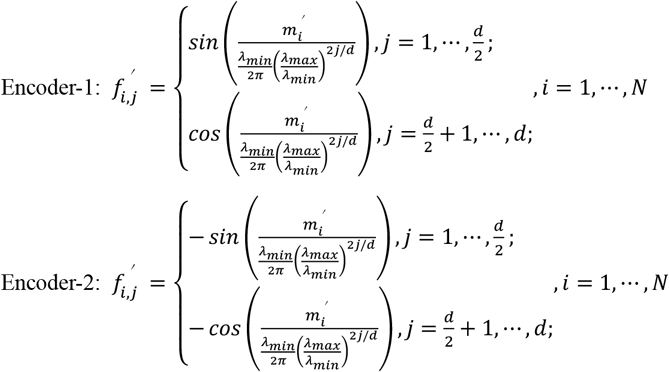

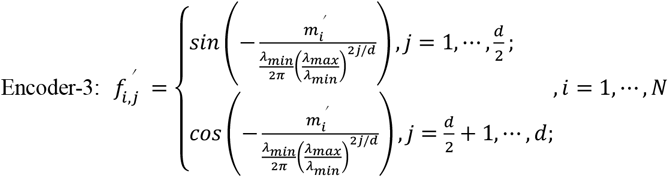

where 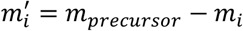, *m*_*precursor*_ represents the mass of the precursor. Each intensity value of the experimental and complementary spectrum is also encoded as a d-dimensional vector. The embedding of the i-th intensity value (*e*_*i*_) is calculated as:

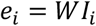

where W is a d-dimensional vector trained during the training phase. Next, the resulting *f*_*i,j*_ and *e*_*i*_ are added together to obtain the output of Peak Encoder E, while 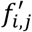 and *e*_*i*_ are added together to obtain the output of Peak Encoder C.

#### Precursor Encoder

A precursor is composed of a mass value (*m*_*prec*_) and a charge value (*c*_*prec*_ ∈ {1,2, ⋯, 10}). The *m*_*prec*_ is embedded using the same sinusoidal embedding as described in Peak Encoder E, while the *c*_*prec*_ value is embedded using an embedding layer. These two embeddings are summed to form the output of the Precursor Encoder.

#### AA Encoder and Positional Encoder

Amino acids are embedded using an embedding layer and then processed by absolute positional embedding^[16]^. These two embeddings are summed together as the input of the Transformer Decoder.

#### Transformer Encoder and Decoder

*π*-HelixNovo consists of a total of *n*_*l*_ Transformer encoder layers with *n*_*h*_ heads that form the Transformer Encoder, and *n*_*l*_ Transformer decoder layers with *n*_*h*_ heads that form the Transformer Decoder. The dimension of *π*-HelixNovo is *d*_*m*_, and the dimension of the feedforward module is *d*_*f*_. Dropout is not used in *π*-HelixNovo.

#### Linear and Softmax modules

The Linear and Softmax module takes the outputs of the Transformer Decoder as input. It produces the probabilities of all possible amino acids and the end symbol corresponding to each position in the peptide. These probabilities are then used to generate the peptide sequence through search strategies.

#### Search strategy

*π*-HelixNovo employs two search strategies, namely greedy search (GS)^[17]^ and beam search (BS)^[18]^, to generate amino acids from the output of the Linear and Softmax modules. Greedy search selects the best solution among the current outputs, while beam search searches in the last *n*_*beams*_ outputs, where *n*_*beams*_ is a hyperparameter. Therefore, the greedy search obtains a local optimal solution, while the beam search obtains a solution between local and global optima. Greedy search is fast but may not produce the most accurate results, while beam search is more accurate but requires more time as *n*_*beams*_ increases. It’s worth noting that when *n*_*beams*_ is set to 1, beam search is equivalent to greedy search. Therefore, the choice of the search strategy in *π*-HelixNovo depends on the trade-off between performance and computational cost.

#### Training hyperparameters

*π*-HelixNovo has around 47 million parameters, as *n*_*l*_, *n*_*h*_, *d*_*m*_, and *d*_*f*_ are specified as 9, 8, 512, and 1024, respectively. The primary training objective is to minimise the cross-entropy (CE) loss^[19]^ between the ground truth amino acid sequence (*R*^*b*×*l*^) and the predicted probabilities (*R*^*b*×*l*×*k*^) outputted by the Linear and Softmax modules. During training, an Adam optimiser with a 10^−5^ weight decay and a cosine learning rate scheduler with 100k warm-up updates are used. Here, *b* (*l*2*m* represents the batch size, *l* is the length of the peptide sequence, and *k* is the number of possible amino acids. To clarify, the models trained on the nine-species benchmark dataset and MSV000081142 dataset involve 20 amino acids and 7 modified amino acids, so the value of *k* is set to 27 for these models. Similarly, *k* is 25 for the model trained on the merged dataset.

#### Identification of novel members of gut microbes

We have conducted a combined database search and de novo sequencing approach for identifying novel peptides in gut microbes. Conceptually, the following procedure can be applied to identify new proteins/peptides in any relevant biological applications. Firstly, we apply the database search method to an MS2 dataset obtained from a proteomics experiment. Secondly, this dataset is divided into two subsets: a set of peptide-spectrum matches (PSMs) and a set of MS2 spectra that remain unidentified. The PSMs are further divided into training and validation sets using a segmentation method. Subsequently, de novo sequencing models are trained on the training set and evaluated on the validation set to identify the best-performing models. Thereafter, de novo sequencing is performed on the set of MS2 spectra that were initially unidentified. This process generates corresponding peptides based on the de novo sequencing results, which need to be confirmed by genomics data or any other relevant form of data.

The de novo sequenced peptides undergo quality control using the T\D\DS strategy based on a target-decoy search approach^[20]^. This strategy involves comparing the peptides against a target database plus a decoy database and database search results, including target PSMs filtered at an experiment-wide FDR < 1% at the peptide level. The target database is constructed using the corresponding bacterial genomes, while the decoy database is generated by combining common protein contaminants and reversed entries of the corresponding genome pools. The generation process replicates the trypsin digestion process, which is a commonly employed proteolytic enzyme used in proteomics research. It should be noted that the peptides in the target database must have a length between 7 and 40 and end with the amino acid lysine or arginine.

In this study, the genomes of strains in a defined model community are concatenated with major dietary protein sequences to represent the corresponding genomes in the MSV000082287 dataset^[15]^. Initially, a 5-fold cross-validation approach is employed using the PSMs from the MSV000082287 dataset to assess the performance of Casanovo and *π*-HelixNovo. The best-performing model is then selected from each approach to perform de novo sequencing on the unidentified MS2 spectra. Following the de novo sequencing, a quality control method is applied to filter out peptides with high confidence. The performance of Casanovo and *π*-HelixNovo is evaluated based on the peptide count and peptide species count. Additionally, taxonomic identification of tryptic peptides is conducted using the Unipept tool (https://unipept.ugent.be/mpa) ^[14]^ to further assess the performance of Casanovo and *π*-HelixNovo.

#### Evaluation metrics

*π*-HelixNovo employs the same evaluation metrics as other de novo sequencing models: amino acid precision (AA precision), amino acid recall (AA recall), and peptide recall. To calculate these metrics, we define the matched amino acid predictions 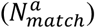 as all predicted amino acids that meet the following condition:

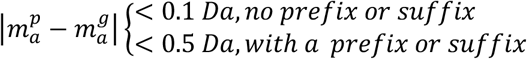

where 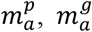 denote the masses of the predicted and ground truth amino acids, respectively. Based on this, we can define the evaluation metrics as follows:

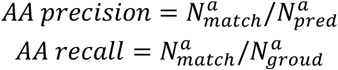

where 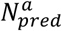 represents the number of predicted amino acids, and 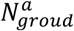 is the number of amino acids in ground truth peptides. Additionally, peptide recall is defined as:

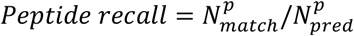

where 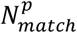 represents the number of matched peptides for which 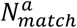 is equal to the length of the peptide.

## Data availability

The nine-species benchmark dataset used in this study is available at https://zenodo.org/record/6791263/files/casanovo_preprocessed_data.zip?download=1, the merged dataset, ABRF, PXD008844, and PXD010559 datasets can be accessed from https://zenodo.org/record/8280267, the MSV000081142 dataset can be downloaded from ftp://massive.ucsd.edu/MSV000081142/peak/filtered_library_mgf_files/ and ftp://massive.ucsd.edu/MSV000081142/updates/2017-12-18_mwang87_194c3d43/peak/filtered_library_mgf_files, which includes a total of 300 mgf files, and the MSV000082287 dataset is available at ftp://massive.ucsd.edu/MSV000082287/.

## Code availability

*To be announced soon*

## Acknowledgements

We would like to acknowledge Beinn Purvis (Kunming Institute of Botany, Chinese Academy of Sciences) for the helpful comments and suggestions. This work has been supported by a direct national funding from Chinese Ministry of Technology to Peng Cheng Laboratory, the National Key Research and Development Program of China (2021YFA1301603 and 2020YFE0202200), Research and Development Program of Guangzhou Laboratory (SRPG22-001), the National Natural Science Foundation of China (32088101), and the CAMS Innovation Fund for Medical Sciences (CIFMS) (2019-I2M-5-063).

## Author Contributions

T.Y. conceived the concept of complementary spectra, designed the architecture of *π*-HelixNovo, performed the experiments, and wrote the initial manuscript. T.L., Z.L., F.X. and X.H. collected the data and performed the data analysis. L.X. helped to improve the model. B.S. and L.L. performed the metaproteomics data analysis. Y.H. and F.H. coordinated the study. Y.W. and C.C. designed the study, analyzed the results, and co-supervised the project. All authors helped to revise the manuscript and approved the final manuscript.

## Competing Interests

The authors have declared they have no conflict of interest.

### Abbreviations

MS: mass spectrum
DDA: data-dependent acquisition
CID: collision-induced dissociation
HCD: higher-energy C-trap dissociation
PSM: peptide-spectrum match
AA: amino acids
GS: greedy search
BS: beam search
CE: cross-entropy.

## Supplementary information

**Supplementary Fig. 1.**
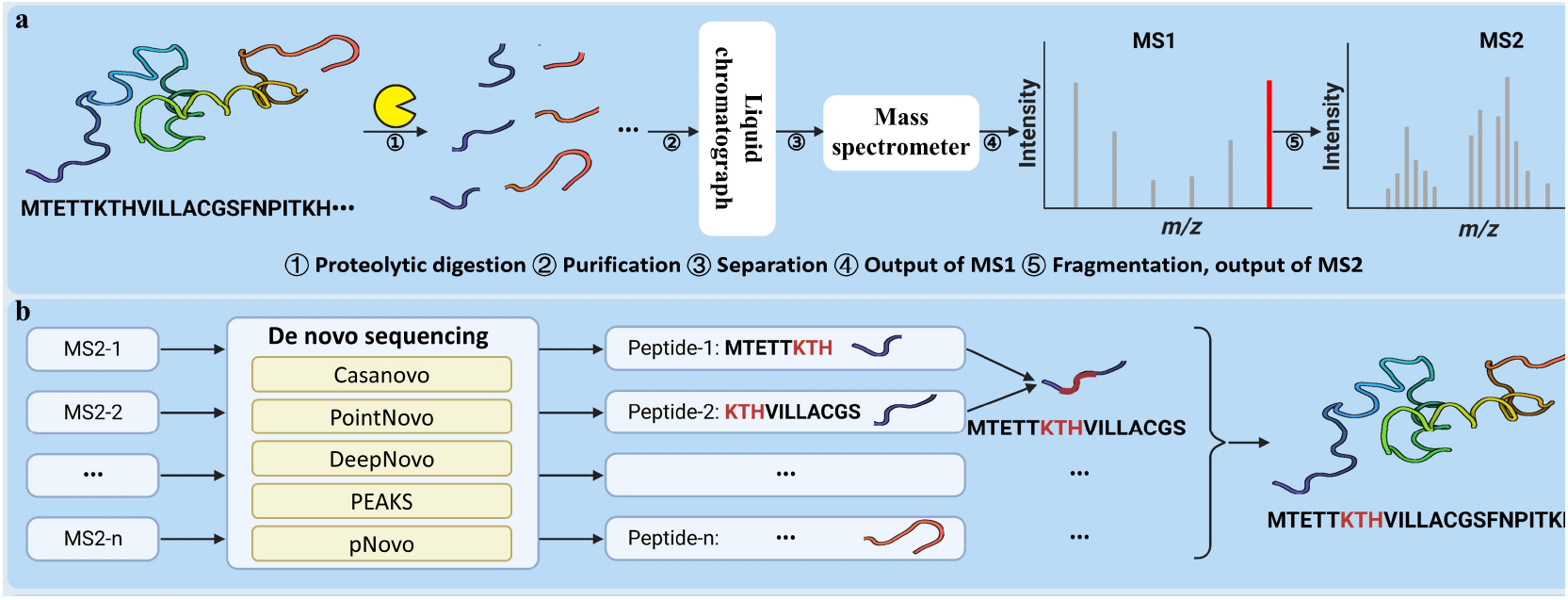
The overview of tandem mass spectrometry and de novo sequencing. (a) The overview of tandem mass spectrometry. (b) The de novo sequencing methods for protein sequence identification tasks and the method of assembling the whole protein sequence from the identified peptides.

**Supplementary Fig. 2.**
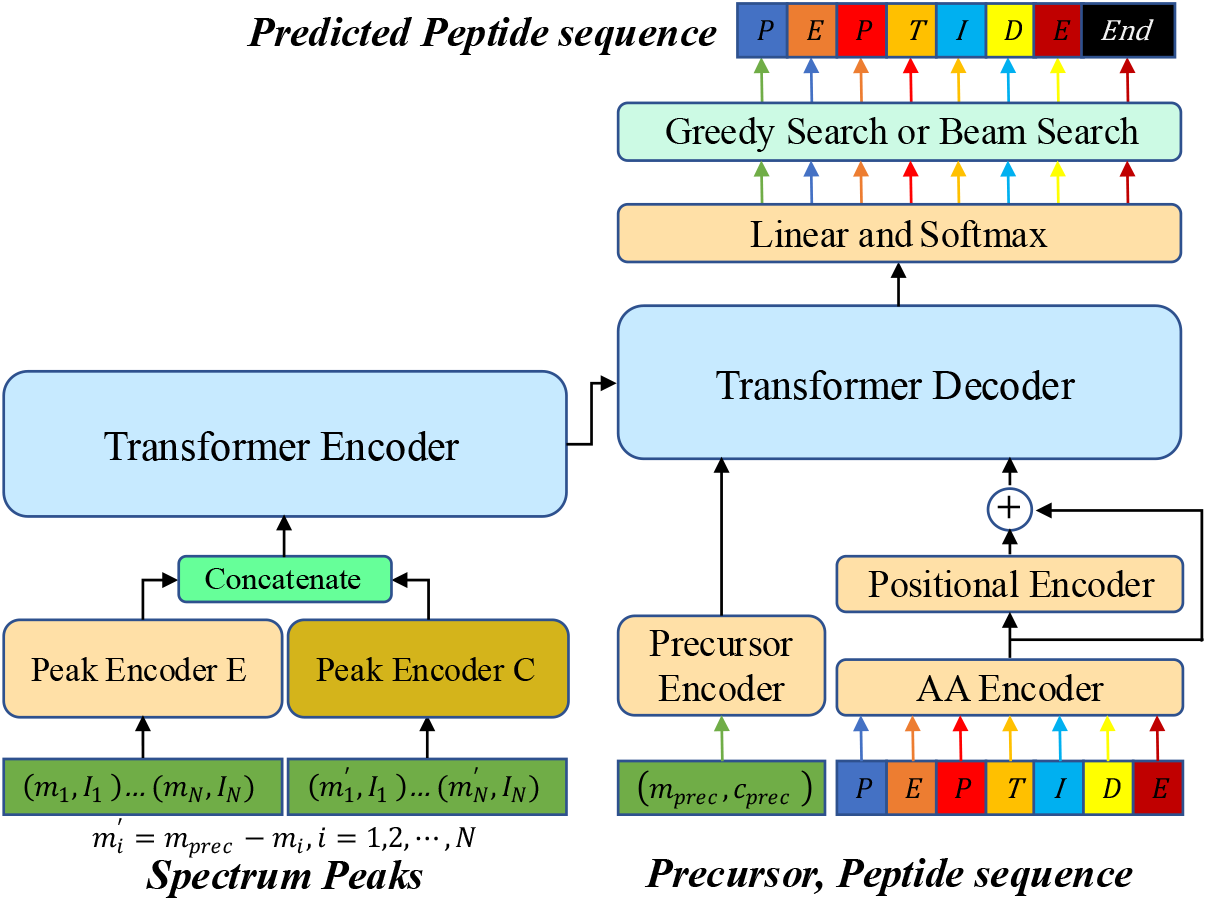
The architecture of *π*-HelixNovo.

**Supplementary Fig. 3.**
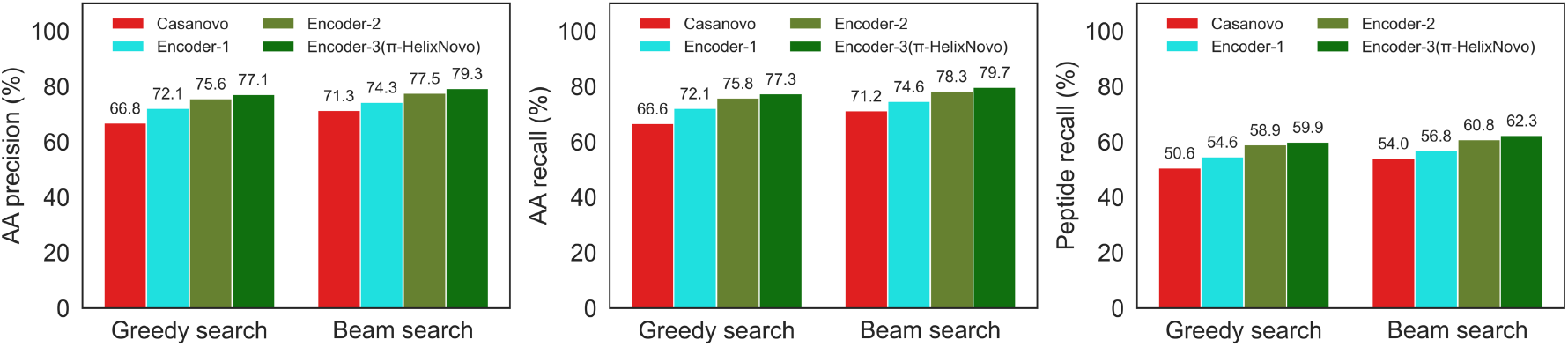
The performance of the models using three different encoders for the complementary spectrum. Models are evaluated using greedy search and beam search (*n*_*beams*_ *=* 2) on *V. mungo* dataset.

**Supplementary Fig. 4.**
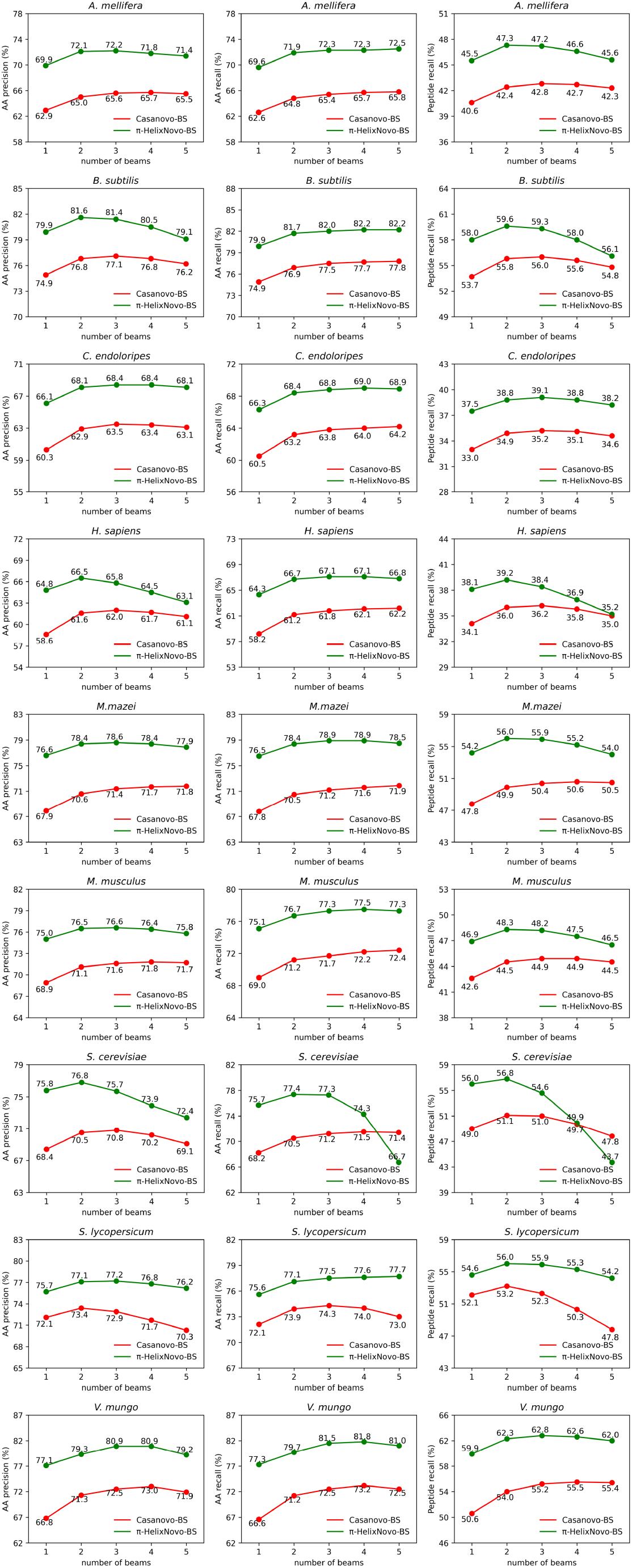
The performance of *π*-HelixNovo and Casanovo using the beam search strategy with different *n*_*beams*_ on the nine-species benchmark dataset.

**Supplementary Fig. 5.**
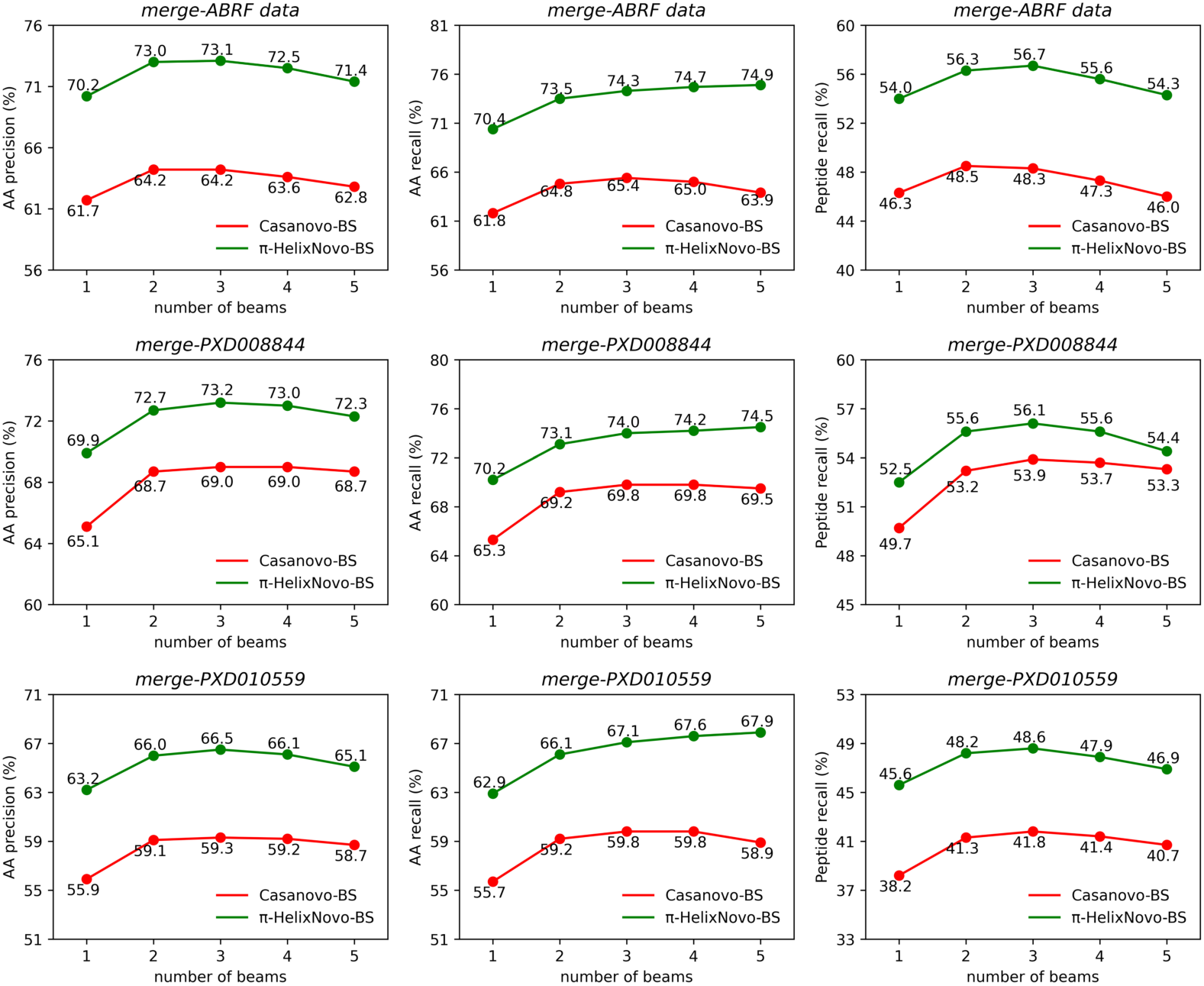
The performance of *π*-HelixNovo and Casanovo trained on the merged dataset and tested on ABRF, PXD008844, and PXD010559 datasets using the beam search strategy with different *n*_*beams*_.

**Supplementary Fig. 6.**
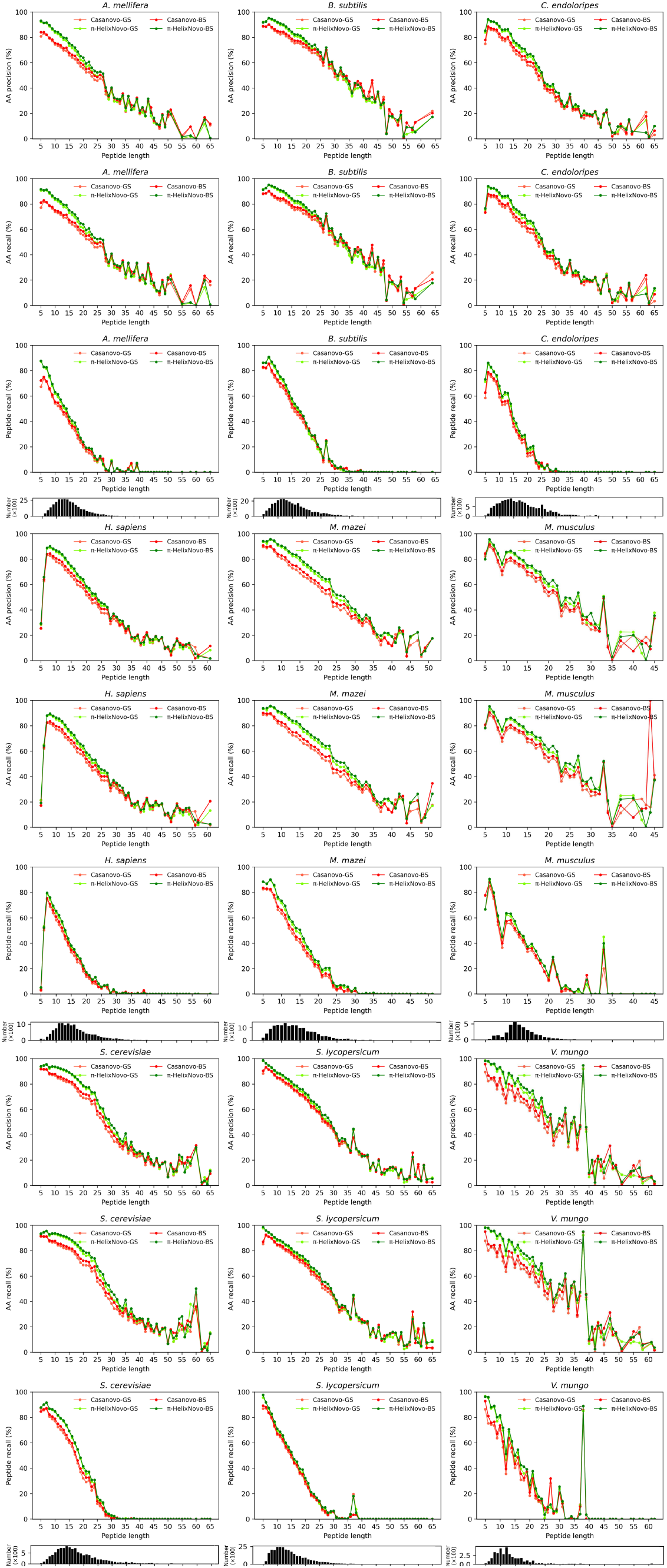
The performance of *π*-HelixNovo and Casanovo on the nine-species benchmark dataset using greedy search and beam search (*n*_*beams* = 2). MS2 spectra with different peptide lengths are evaluated, and the distribution map of peptide length is shown below (Y- axis is the number of peptides, X-axis is the peptide length).

**Supplementary Fig. 7.**
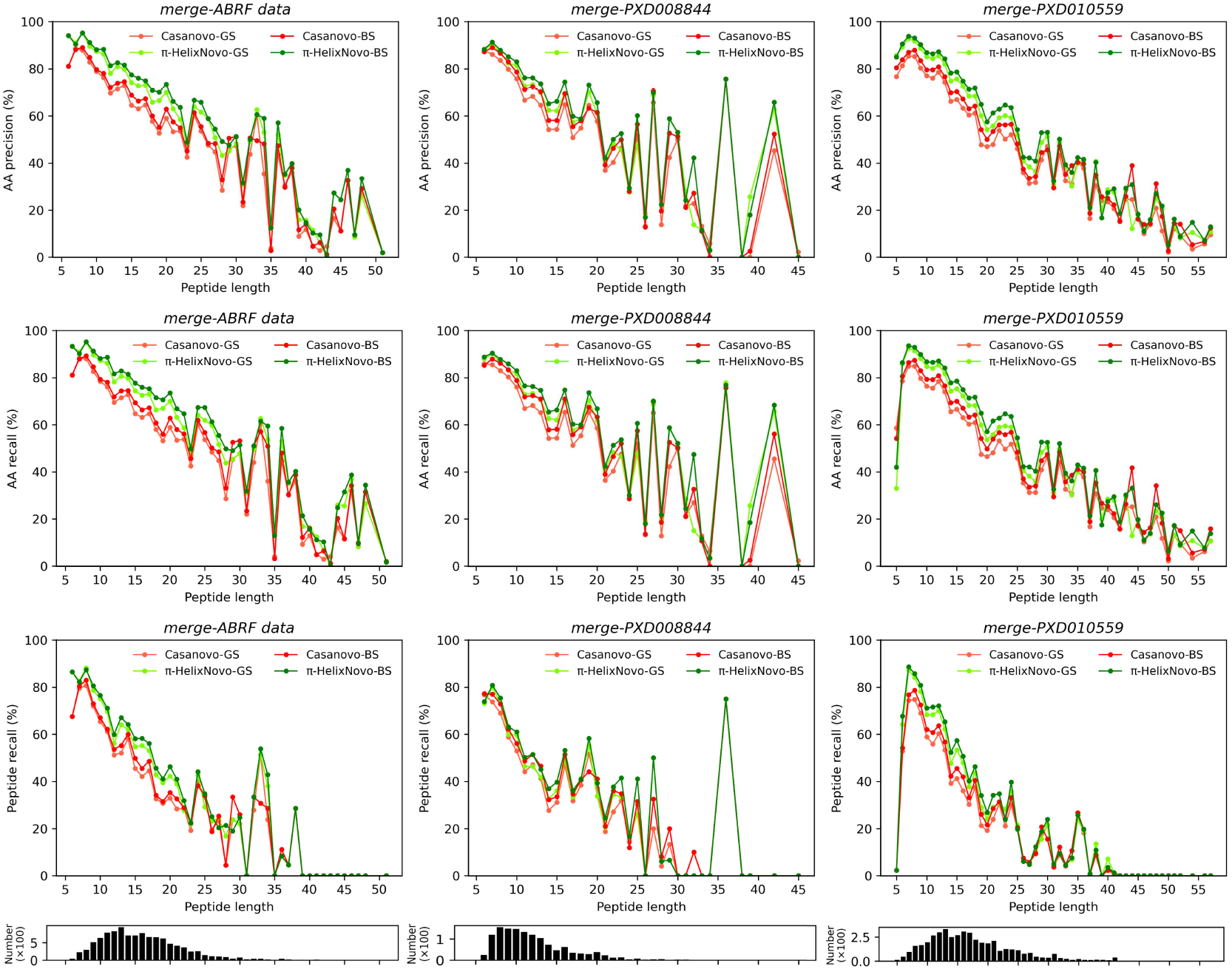
The performance of *π*-HelixNovo and Casanovo trained on the merged dataset. MS2 with different peptide lengths in ABRF, PXD008844, and PXD010559 datasets are evaluated using greedy search and beam search (*n*_*beams* = 2), and the distribution map of peptide length is shown below (the number of peptides is on the Y-axis, peptide length is on the X-axis).

**Supplementary Fig. 8.**
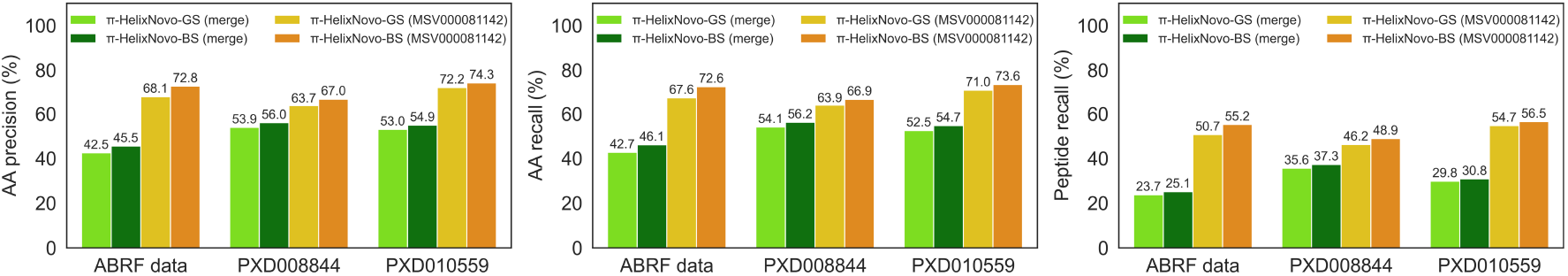
The performance of *π*-HelixNovo on the three processed test datasets in which the peptide sequences are never before seen. Merge means *π*-HelixNovo was trained on the merged dataset and MSV000081142 means *π*-HelixNovo was trained on the MSV000081142 dataset

**Supplementary Table 1.** The time cost of *π*-HelixNovo-BS on the *M. mazei* dataset (164412 spectra).

| Supplementary Table 1\| The time cost of π-HelixNovo-BS on the *M. mazei* dataset (164412 spectra). | | | | | |
| --- | --- | --- | --- | --- | --- |
| $n_{beams}$ | 1 | 2 | 3 | 4 | 5 |
| time cost | 40 min | 12 h 30 min | 24 h | 33 h 22 min | 42h 40min |

**Supplementary Table 2.**
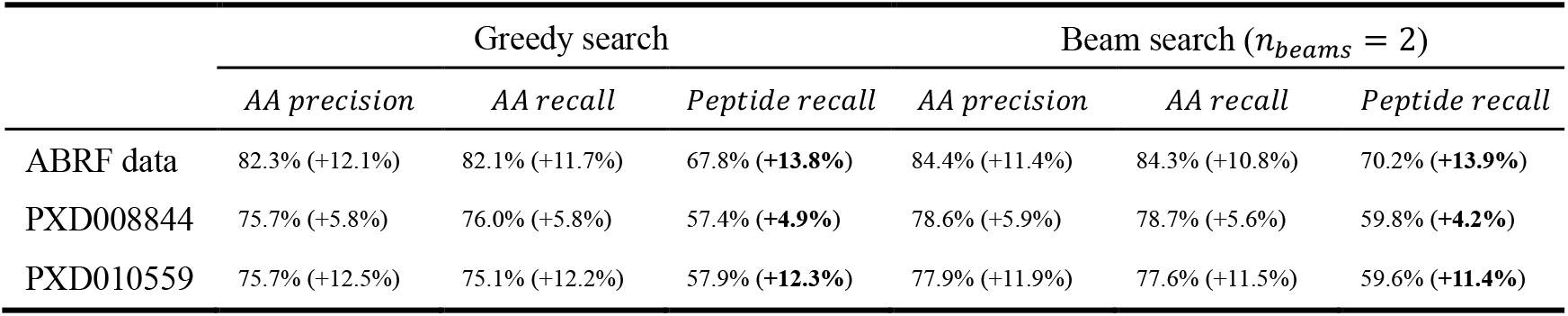
The performance of *π*-HelixNovo trained on the MSV000081142 dataset, compared to *π*-HelixNovo trained on the merged dataset.

| Supplementary Table 2\| The performance of π-HelixNovo trained on the MSV000081142 dataset, compared to π-HelixNovo trained on the merged dataset. | | | | | | |
| --- | --- | --- | --- | --- | --- | --- |
| | Greedy search | | | Beam search ($n_{beams} = 2$) | | |
|  | *AA precision* | *AA recall* | *Peptide recall* | *AA precision* | *AA recall* | *Peptide recall* |
| ABRF data | 82.3% (+12.1%) | 82.1% (+11.7%) | 67.8% (+13.8%) | 84.4% (+11.4%) | 84.3% (+10.8%) | 70.2% (+13.9%) |
| PXD008844 | 75.7% (+5.8%) | 76.0% (+5.8%) | 57.4% (+4.9%) | 78.6% (+5.9%) | 78.7% (+5.6%) | 59.8% (+4.2%) |
| PXD010559 | 75.7% (+12.5%) | 75.1% (+12.2%) | 57.9% (+12.3%) | 77.9% (+11.9%) | 77.6% (+11.5%) | 59.6% (+11.4%) |

**Supplementary Table 3.**
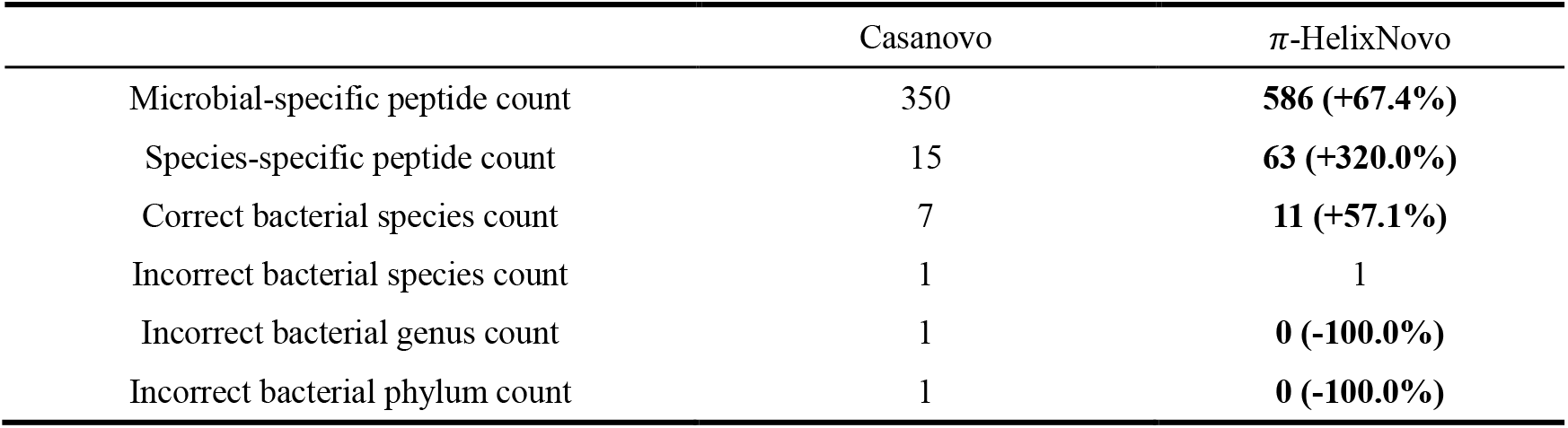
The analysis of the peptides through QC (T\D\DS) for Casanovo and *π*-HelixNovo. T\D\DS means a peptide was identified in target database but absent in decoy database and database search results.

